# Emergence of *optrA*-mediated linezolid resistance in multiple lineages and plasmids of *Enterococcus faecalis* revealed by long read sequencing

**DOI:** 10.1101/2020.02.28.969568

**Authors:** Martin P McHugh, Benjamin J Parcell, Kerry A Pettigrew, Geoff Toner, Elham Khatamzas, Anne Marie Karcher, Joanna Walker, Robert Weir, Danièle Meunier, Katie L Hopkins, Neil Woodford, Kate E Templeton, Stephen H Gillespie, Matthew TG Holden

## Abstract

**Objectives:** To characterise the genetic environment of *optrA* in linezolid-resistant *Enterococcus faecalis* isolates from Scotland.

**Methods:** Linezolid-resistant *E. faecalis* were identified in three Scottish Health Boards and confirmed to carry the *optrA* gene at the national reference laboratory. WGS was performed with short read (Illumina MiSeq) and long read (Oxford Nanopore MinION) technologies to generate complete genome assemblies. Illumina reads for 94 *E. faecalis* bloodstream isolates were used to place the *optrA*-positive isolates in a larger UK phylogeny.

**Results:** Six *optrA*-positive linezolid-resistant *E. faecalis* were isolated from urogenital samples in three Scottish Health Boards (2014-2017). No epidemiological links were identified between the patients, four were community-based, and only one had recent linezolid exposure. Reference-based mapping confirmed the isolates were genetically distinct (>13,900 core SNPs). *optrA* was located on a plasmid in each isolate and these plasmids showed limited nucleotide similarity. There was variable presence of transposable elements surrounding *optrA*, (including IS*1216*, IS*3*, and Tn*3*) and not always as a recognisable gene cassette. OptrA amino acid sequences were also divergent, resulting in four protein variants differing in 1-20 residues. One isolate belonged to ST16 and clustered with three other isolates in the UK collection (76-182 SNPs), otherwise the *optrA-*positive isolates were genetically distinct from the bloodstream isolates (>6,000 SNPs).

**Conclusions:** We report multiple variants of the linezolid resistance gene *optrA* in diverse *E. faecalis* strain and plasmid backgrounds, suggesting multiple introductions of the gene into the *E. faecalis* population and selection driving recent emergence.

## INTRODUCTION

*Enterococcus faecalis* and *Enterococcus faecium* are carried in the intestinal tract and are important opportunistic pathogens in humans.^1^ Treatment of enterococcal infections is challenging due to intrinsic or acquired resistance to multiple antimicrobials including aminoglycosides, benzylpenicillin, cephalosporins, fluoroquinolones, macrolides, tetracyclines, and trimethoprim. Among the remaining treatment options, clinical *E. faecium* isolates are usually resistant to amoxicillin and resistance to vancomycin is increasingly common.^2^ In contrast, *E. faecalis* typically remains susceptible to amoxicillin and vancomycin but can acquire significant resistance and has been implicated in the transfer of antimicrobial resistance genes to other Gram-positive pathogens, for example transmitting *vanA*-mediated vancomycin resistance to methicillin-resistant *Staphylococcus aureus*.^3^

The main treatment options for multi-drug resistant Gram-positive bacteria are the oxazolidinones linezolid or tedizolid, or the lipopeptide daptomycin. Daptomycin therapy is challenging due to significant side effects, limited efficacy in pulmonary infections, uncertain dosing regimens, and challenges with *in vitro* susceptibility determination.^4^ Linezolid blocks protein synthesis by binding to the 50S ribosomal subunit and inhibiting formation of the initiation complex.^5^ Linezolid resistance is uncommon, reported in ≤1% of bloodstream enterococcal isolates in the UK.^6,7^ The G2576T mutation in the 23S rRNA genes can arise *de novo* during extended linezolid therapy,^8^ although strict infection control and antimicrobial stewardship have been successful in limiting incidence.^9^ The methyltransferases Cfr and Cfr(B), and ABC-F ribosomal protection proteins OptrA and PoxtA also confer resistance to linezolid but are carried on mobile genetic elements, raising the prospect of rapid spread of linezolid resistance across genetically distinct lineages.^10–12^ In 2015, *optrA* was first reported as conferring resistance to oxazolidinones and phenicols.^13^ Recent international surveillance shows that although linezolid resistance remains rare, *optrA* has spread to every continent and is the dominant mechanism of linezolid resistance in *E. faecalis*.^14^ Surveillance has also detected *optrA* in the UK.^15^ Studies into the genetic context of *optrA* have identified the gene on both the chromosome and plasmids, often associated with insertion sequence IS*1216*, a possible explanation for the rapid spread of *optrA*.^16,17^ However, few studies have generated complete genome assemblies of *optrA-*carrying *E. faecalis*, which would provide high precision information on the genetic context of *optrA*.

Here, we investigate the epidemiological and clinical background of *optrA*-carrying *E. faecalis* isolates from human clinical samples collected in Scotland. We used whole genome sequencing to determine whether these isolates represent transmission of a single clonal lineage. We hypothesised the spread of *optrA* is driven by a single mobile genetic element, and to investigate this we made hybrid assemblies of short and long read sequencing data to generate complete genomes and to reconstruct the genetic environment of *optrA*. This study describes the first use of nanopore-based long read sequencing to investigate *optrA*-containing mobile genetic elements.

## MATERIALS AND METHODS

### Bacterial strains

Isolates were selected for this study based on the presence of the *optrA* gene as determined by Public Health England’s Antimicrobial Resistance and Healthcare Associated Infections (AMRHAI) Reference Unit, either as part of non-structured retrospective screening of stored isolates (prior to 2016) or as part of the reference laboratory service (2016 onwards). Isolates were originally collected in three Scottish Health Boards, and as such represent a subset of Scottish *optrA*-positive isolates identified by AMRHAI. Linezolid- and chloramphenicol-resistant *E. faecalis* were isolated from six clinical samples (Table 1) using standard methods and identified with matrix-assisted laser-desorption ionisation time-of-flight mass spectrometry or the Vitek-2 GP-ID card (bioMérieux, Marcy L’Etoile, France). Antimicrobial susceptibility testing was performed with the Vitek-2 AST-607 card and interpreted with EUCAST breakpoints.^18^ Isolates were referred to AMRHAI for characterisation of linezolid resistance mechanisms. Detection of the G2576T mutation (*Escherichia coli* numbering) in the 23S rRNA genes was investigated by PCR-RFLP and, from 2016, by a real-time PCR-based allelic discrimination assay.^19,20^ The *cfr* and *optrA* genes were sought by a multiplex PCR using primers for the detection of *cfr* (*cfr-fw*: 5’-TGA AGT ATA AAG CAG GTT GGG AGT CA-3’ and *cfr-rev:* 5’-ACC ATA TAA TTG ACC ACA AGC AGC-3’)^21^ and for the detection of *optrA* (*optrA-F*: 5’-GAC CGG TGT CCT CTT TGT CA-3’ and *optrA-R*: 5’-TCA ATG GAG TTA CGA TCG CCT-3’) (AMRHAI, unpublished data).

**Table 1.**
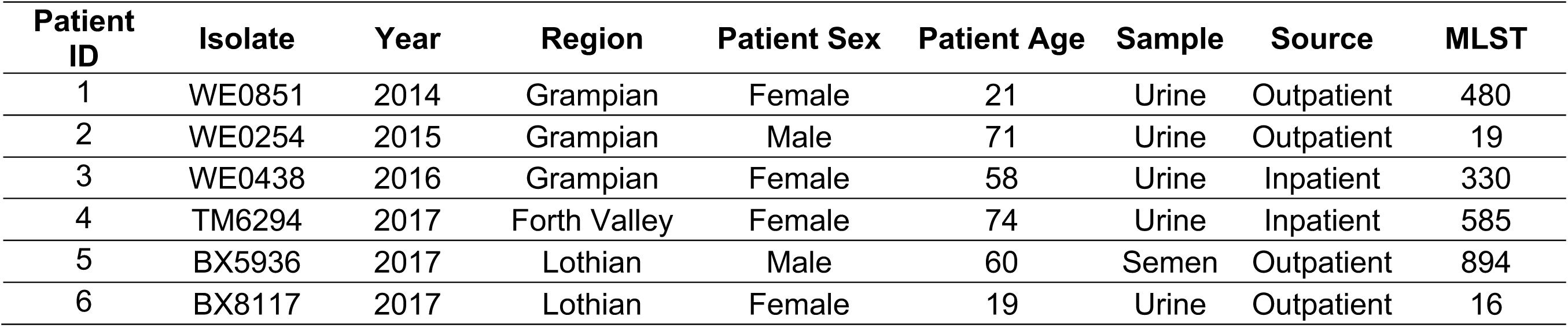
Details of the *optrA*-positive *E. faecalis* characterized in this study

| **Patient ID** | **Isolate** | **Year** | **Region** | **Patient Sex** | **Patient Age** | **Sample** | **Source** | **MLST** |
| --- | --- | --- | --- | --- | --- | --- | --- | --- |
| 1 | WE0851 | 2014 | Grampian | Female | 21 | Urine | Outpatient | 480 |
| 2 | WE0254 | 2015 | Grampian | Male | 71 | Urine | Outpatient | 19 |
| 3 | WE0438 | 2016 | Grampian | Female | 58 | Urine | Inpatient | 330 |
| 4 | TM6294 | 2017 | Forth Valley | Female | 74 | Urine | Inpatient | 585 |
| 5 | BX5936 | 2017 | Lothian | Male | 60 | Semen | Outpatient | 894 |
| 6 | BX8117 | 2017 | Lothian | Female | 19 | Urine | Outpatient | 16 |

An *E. coli* transformant harbouring a plasmid bearing *cfr* (kindly provided by Pr S. Schwarz) was used as a control strain for the detection of *cfr*. This was replaced from 2016 by *Staphylococcus epidermidis* NCTC 13924 harbouring both *cfr* and the G2576T mutation. *E. faecium* NCTC 13923 was used as a control strain for the detection of *optrA*.

Access to isolates and clinical data was approved by the NHS Scotland Biorepository Network (Ref TR000126).

### Whole genome sequencing and genomic analysis

Single colonies were inoculated into brain heart infusion broth (Oxoid, Basingstoke, UK) and incubated overnight at 37°C. Genomic DNA was extracted from cell pellets using the Wizard Genomic DNA Purification Kit (Promega, Wisconsin, USA), or QiaSymphony DSP DNA Mini Kit (Qiagen, Hilden, Germany). Short read barcoded libraries were prepared using the Nextera XT kit (Illumina, San Diego, USA) and sequenced with a MiSeq instrument (Illumina) using 250 bp paired-end reads on a 500-cycle v2 kit. Short reads were quality trimmed with Trimmomatic v0.36 and settings [LEADING:5 TRAILING:5 SLIDINGWINDOW:4:15 MINLEN:100].^22^ Barcoded long read libraries were generated with the 1D Ligation Sequencing Kit (Oxford Nanopore Technologies, Oxford, UK) and sequenced with an R9.4 flow cell on a MinION sequencer (Oxford Nanopore Technologies). Base-calling and barcode de-multiplexing was performed with Albacore v2.1.3 (Oxford Nanopore Technologies) and the resulting fast5 files converted to fastq with Poretools v0.6.0,^23^ or basecalled and de-multiplexed with Albacore v2.3.3 with direct fastq output. Porechop v0.2.3 (https://github.com/rrwick/Porechop) was used to remove chimeric reads and trim adapter sequences. The data for this study have been deposited in the European Nucleotide Archive (ENA) at EMBL-EBI under accession number PRJEB36950 (https://www.ebi.ac.uk/ena/data/view/PRJEB36950).

To generate a UK-wide *E. faecalis* phylogenetic context for the *optrA*-positive isolates, raw sequence data was downloaded from the ENA (www.ebi.ac.uk/ena) under study accession numbers PRJEB4344, PRJEB4345, and PRJEB4346.^24^ Short reads were mapped to the *E. faecalis* reference genome V583 (accession number AE016830) using SMALT v0.7.4.^25^ Mapped assemblies were aligned and regions annotated as mobile genetic elements in the V583 genome (transposons, integrases, plasmids, phages, insertion sequences, resolvases, and recombinases; tab file of regions available in Table S1) were removed from the assembly (https://github.com/sanger-pathogens/remove_blocks_from_aln). All sites in the alignment with single nucleotide polymorphisms (SNPs) were extracted using SNP-sites v2.4.0^26^ and a phylogeny was created from the core-genome SNP alignment using RAxML v8.2.8^27^ with 100 bootstrap replicates and visualised with iTOL.^28^ Recombination blocks were removed from ST16 isolates using Gubbins v1.4.10.^29^

Hybrid assembly was performed with Illumina short reads and Nanopore long reads using Unicycler v0.4.7^30^ in standard mode. The resulting assemblies were annotated with Prokka v1.5.1 using a genus specific RefSeq database.^31^ Hybrid assemblies were checked for indel errors using Ideel (https://github.com/mw55309/ideel) and UniProtKB TrEMBL database v2019_1. Plasmid comparisons were generated and visualised with EasyFig v2.2.2^32^ and BRIG v0.95.^33^

MLST typing was performed using SRST2 v0.2.0^34^ and the *E. faecalis* MLST database (https://pubmlst.org/efaecalis/) sited at the University of Oxford.^35,36^ Antimicrobial resistance mechanisms were detected using ARIBA v2.12.1^37^ and the ResFinder database v3.0^38^ with the addition of linezolid resistance mutations in the 23S rRNA (G2505A and G2576T based on *E. coli* numbering).

## RESULTS AND DISCUSSION

### Detection of *optrA*-positive E. faecalis

Six *E. faecalis* isolated from urogenital samples were initially identified as linezolid- and chloramphenicol-resistant in routine diagnostic laboratories and confirmed to carry *optrA* at the AMRHAI Reference Unit (Table 1). The earliest isolates in this collection were from the Grampian region of Scotland in 2014, 2015, and 2016. Three more isolates were identified in 2017 from other regions of Scotland (Forth Valley and Lothian, Table 1), with no clear epidemiological links between the patients. Prior to isolation of *optrA-*positive *E. faecalis*, patients 1-3 were treated with trimethoprim for recurrent urinary tract infections, with patient 3 also receiving cefalexin. Patients 5 and 6 were managed in general practice and it was not possible to determine their antimicrobial exposure. Patient 4 was the only patient with known exposure to linezolid, a two-week course prior to the isolation of *optrA-*positive *E. faecalis*. Patient 4 was a surgical inpatient and another patient on the same ward had *optrA-*positive *E. faecalis* isolated from an abdominal wound, indicating possible transmission between this patient and patient 4. The *optrA*-positive *E. faecalis* from the contact of patient 4 was not available for study. Further screening of the ward environment and patients found no further linezolid-resistant enterococci or staphylococci over a two-month period, although the contact of patient 4 continued to have *optrA*-positive *E. faecalis* isolated from their abdominal wound for a month until discharge.

### *optrA* is carried by distinct strains

Whole genome sequencing was performed to investigate the genetic relationship between the isolates. *In silico* MLST showed the six isolates belonged to different sequence types (STs), suggesting they were genetically distinct (Table 1). To further confirm this, we analysed SNPs in the core genomes of the *optrA*-positive isolates and found the isolates differed by a median 18,806 SNPs (range 13,909 – 22,272). Previous estimates suggest a genetic diversification rate of 2.5-3.4 SNPs/year for *E. faecalis*, highlighting the *optrA*-positive strains share a very distant common ancestor.^24^

### *optrA* is carried on diverse plasmids

We then examined the genetic context of *optrA* in each isolate. Initial *de novo* assembly of short-read data generated fragmented assemblies (72-135 contigs, mean N50 198 kb), but with *optrA* present on moderate sized contigs (11-44 kb). Three *optrA*-positive contigs carried plasmid-associated replication or transfer genes, but none represented a complete plasmid, or had increased read depth coverage compared to core genes indicative of being multicopy. Therefore, it was unclear if *optrA* was carried on plasmids (often present in multiple copies within a cell) or the chromosome, and how similar these regions were between the six isolates. To resolve repetitive regions and try to complete the genome assemblies we utilised Nanopore sequencing to generate long reads, and then combined these with Illumina short reads to produce high quality hybrid assemblies. In four of the isolates completed genomes were obtained, with the other two generating near-complete genomes (Table S2). Analysis of the six hybrid assemblies showed <3 % putative coding sequences were shorter than the closest reference match (Table S2) indicating the hybrid assembly process removed most indel errors and the short coding sequences were likely to be true pseudogenes.^39^ The hybrid assemblies contained between one and three plasmids ranging in size from 11-80 kb, with *optrA* present on a single complete plasmid in each isolate (Table 2).

**Table 2.** Plasmids from Hybrid Assemblies

| Isolate | Element | Copy Number<sup>a</sup> | Size (bp) | Plasmid rep type | Best NCBI match | Resistance genes | Variation compared to *optrA*<sub>pE394<sup>b</sup></sub> | Transposases (n) |
| --- | --- | --- | --- | --- | --- | --- | --- | --- |
| BX5936 | pBX5936-1 | 1 | 68656 | rep9 | Efs pE035, coverage 65%, 98% ID (MK140641) | *fexA*, *optrA* | S2F | ISEf1 (2)<br>IS1216 (2) |
|  | pBX5936-2 | 1 | 51669 | rep9 | Efs FC unnamed plasmid1, coverage 85%, 100% ID (CP028836) | *catA8*, *tet(L)*, *tet(M)*, *ant(6)-la*, *cadA*, *copZ*, *ermB* | - | NA |
| BX8117 | pBX8117-1 | 1 | 68773 | rep9 | Efs FDAARGOS_324 unnamed plasmid2, coverage 100%, 100% ID (CP028284) | None | - | NA |
|  | pBX8117-2 | 1 | 41839 | rep9 | Efs pEF123, coverage 64%, 98% ID (KX579977) | *catA8*, *cfr(D)*, *optrA*, *fexA* | K3E, N12Y, E37K, N122K, Y135C, Y176D, A350V, V395A, A396S, Q509K, Q541E, M552L, N560Y, K562N, Q565K, E614Q, I627L, D633E, N640I, R650G | IS1216 (5) |
| TM6294 | pTM6294-1 | 1 | 75362 | rep9 | Efs FC unnamed plasmid1, coverage 74%, 99% ID (CP028836) | *catA8*, *tet(L)*, *tet(M)*, *ant(6)-la*, *cadA*, *copZ*, *ermB*, *aph(3')-IIIa*, *sat4*, *ant(6)-la*, *lnuB*, *lsaE*, *ant(9)*, *ant(6)-la*, *aac(6')-le-aph(2'')-la, aadK, ermB, dfrG* | - | NA |
|  | pTM6294-2 | 1 | 52776 | rep9 | Efs pE035, coverage 87%, 99% ID (MK140641) | *fexA, optrA* | None | ISL3-family (1)<br>IS1216 (1) |
| WE0254 | pWE0254-1 | 1 | 80496 | repUS11 | Efs FDAARGOS_324 unnamed plasmid3, coverage 49%, 99% ID (CP028283) | *ant(9)-la, ermA-like, fexA, optrA* | None | IS3-family (8)<br>IS1216-partial (1) |
|  | pWE0254-2<sup>c</sup> | 1 | 79293 | NA | Efs NCTC8732 chromosome, coverage 99%, 100% ID (LR594051) | None | - | NA |
| WE0438 | pWE0438 | 1 | 61284 | rep9 | Efs pEF123, coverage 76%, 99% ID (KX579977) | *tet(L), tet(M), bcrA, cadA, copZ, ant(6)-la, optrA, fexA, ermB* | K3E, Y176D, I622M | IS1216 (1)<br>ISEnfa1 (2)<br>IS3-family (4)<br>IS6-family (1)<br>Tn3-family (2) |
| WE0851 | pWE0851-1 | 1 | 59708 | repUS11 | Efs pEF123, coverage 22%, 100% ID (KX579977) | *fexA, optrA, ermA-like* | T112K, Y176D | IS1216 (1)<br>IS3-family (6)<br>Tn3-family (1) |
|  | pWE0851-2 | 1 | 26996 | repUS11 | Efs pKUB3007-3, coverage 63%, 100% ID (AP018546) | *aac(6')-le-aph(2'')-la* | - | NA |
|  | pWE0851-3 | 3 | 10826 | NA | Efs pE035, coverage 63%, 99% ID (MK140641) | *aac(6')-le-aph(2'')-la, aac(6')-le-aph(2'')-la, aadK, ermB, ant(6)-la, aph(3')-IIIa, sat4* | - | NA |

bp, base pairs; Efs, *E. faecalis*; ID, identity; NA, not analysed
a Inferred from depth of coverage relative to chromosomal fragment in hybrid assembly
b Amino acid sequence variants compared to the first described *optrA* sequence from pE394 (KP399637)
c Incompletely assembled, in four contigs

In general, the *optrA*-positive plasmids had limited sequence identity, although pBX5936-1 (69 kb) and pTM6294-2 (53 kb) had 97% average nucleotide identity over 40 kb aligned sequence. These two plasmids also had several unique regions indicating the plasmids shared a backbone but had distinct additional content (Figure S1). *optrA* and the phenicol resistance gene, *fexA*, were located in the same orientation and within 550-750 nucleotides of each other, with a short (∼200 nucleotides) hypothetical coding sequence in the intervening region (Figure 1). Additionally, limited similarity was seen between the Scottish *optrA*-positive plasmids and the first identified *optrA*-positive plasmid from China (pE394, accession KP399637; Figure S2).

**Figure 1.**
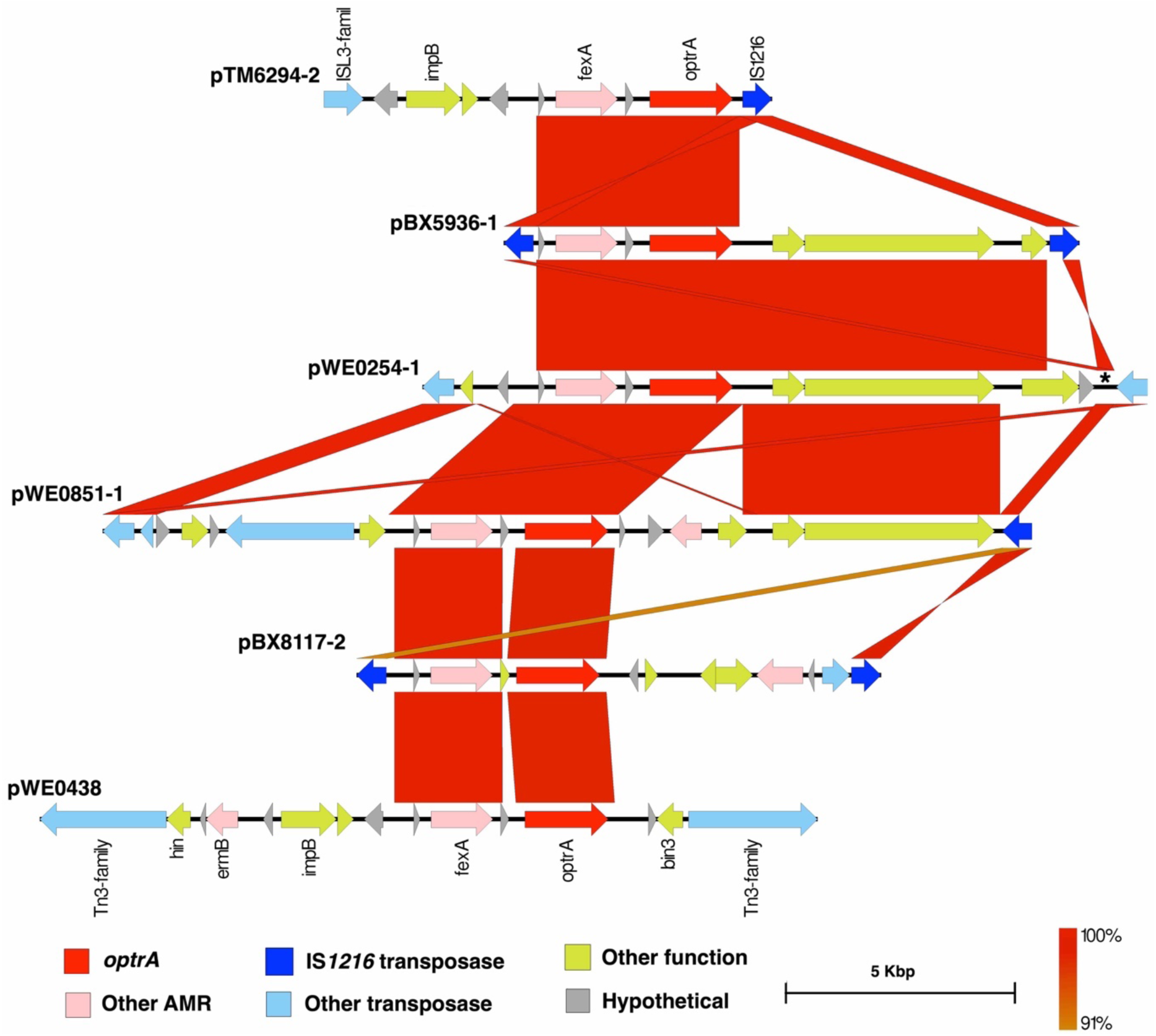
Comparison of *optrA* genetic environments. Alignment of *optrA*-carrying regions shows limited shared sequence identity, apart from around the *optrA* and *fexA* genes. There is evidence of IS*1216* near to the *optrA* gene in all isolates (partial sequence in pWE0254-1 indicated by asterisk), but other insertion sequences are also present suggesting multiple means of *optrA* transmission and ongoing diversification of the element. Arrows indicate coding sequences, blocks between each sequence indicate regions with BLASTn sequence identity >90% and length >100bp.

A number of insertion sequence transposases were identified in the *optrA*-positive plasmids, although we were unable to identify many beyond the family level due to limited matches in public databases (Table 2). We found evidence of IS*1216* in all the *optrA*-positive plasmids, although only pBX5936-1 and pBX8117-2 had IS*1216* flanking the *optrA* and *fexA* region as a cassette (Figure 1). BLASTn comparison of pWE0254-1 with the other *optrA*-positive plasmids highlighted a partial IS*1216* transposase that was not identified by automated annotation. Immediately upstream of the partial IS*1216* was an IS*3*-family transposase, the insertion of which likely disrupted the IS*1216* (Figure S1). pWE0438 had Tn*3*-family transposases surrounding *optrA* and *fexA*, as well genes encoding resolvases, which may represent a transposable unit (Figure 1). pWE0851-1 carried one IS*1216* transposase upstream and one Tn*3*-transposase downstream of *optrA*/*fexA*, but also had multiple copies of IS*3*-family transposases throughout the plasmid so multiple possible mechanisms of transposition exist (Table 2, Figure S1). pTM6294-2 had one IS*1216* transposase and one IS*L3*-family transposase surrounding *optrA*/*fexA*. The variable presence of IS*1216* in these isolates suggest other means of transposition may also be important in the spread of *optrA*, including IS*3*-family and Tn*3*-family transposases.

### *optrA* sequences vary between isolates

Comparison of the OptrA amino acid sequence from each isolate revealed different variants of the resistance protein: two isolates had the same sequence as the first identified OptrA from pE394, BX5936 had a single substitution, WE0851 had two substitutions, WE0348 had three substitutions, and BX8117 had 20 substitutions (Table 2). BX5936 and BX8117 had novel OptrA sequences not yet described in the literature. The OptrA sequence from BX8117 was similar to E35048 detected in an *E. faecium* isolated in Italy in 2015 with the two OptrA sequences differing at three amino acid positions.^40^ Over 40 OptrA sequence variants have been described, although the role of this sequence variation is unclear as they do not significantly differ in their linezolid minimum inhibitory concentration *in vitro*.^41,42^

The degree of sequence variation between the six FexA proteins was less than that seen in OptrA. Comparison to the first reported FexA (AJ549214) showed four common variants in all strains (A34S, L39S, I131V, and V305I), with all but BX8117 having an additional D50A variant. This suggests there is a more diverse background of *optrA* sequences compared to *fexA*, and/or there is ongoing diversifying selective pressure applied only to *optrA* despite the close genetic linkage of the two genes.

Of note, all six isolates had an inferred OptrA sequence 18 amino acids shorter than most public sequences. Inclusion of the 18 upstream amino acids showed that all six isolates had an M1L variant compared to OptrA_pE394_. We believe this is an artefact introduced during coding sequence prediction. The first *optrA* genes were identified *in silico* using ORFfinder (https://www.ncbi.nlm.nih.gov/orffinder), which detects putative coding sequences based on the presence of in-frame start and stop codons only. We used Prokka for genome annotation which implements Prodigal to score potential coding sequences based on start/stop codon position, coding sequence length, and upstream promoter regions and outputs the highest confidence coding sequences. Indeed, most published *optrA* sequences start with nucleotide codons TTG (usually encoding leucine), but the corresponding amino acid sequences start with methionine indicating this codon has been designated as a start codon. The only other report of the M1L variant is from a study that also used Prokka for annotation.^43^ Given the possible effect of methodology on identification of the first amino acid we have not reported the M1L variant in our results but mention it here for completeness. The true *optrA* start codon should be confirmed to aid ongoing surveillance efforts.

### *optrA*-positive strains are distantly related to bloodstream isolates

To investigate whether the *optrA*-positive isolates represented common *E. faecalis* strains in the UK, publicly available sequence data of 94 *E. faecalis* isolates from the British Society for Antimicrobial Chemotherapy (BSAC) bacteraemia surveillance programme (isolated between 2001 and 2011) were analysed together with the six known *optrA*-positive isolates.^24^ We first looked for determinants of linezolid resistance in the 94 sequences, and found no evidence of *cfr, cfr*(B), *cfr*(D), *optrA, poxtA*, or the G2505A 23S rRNA gene mutation. Only one of the BSAC isolates (accession ERS324700) carried the G2576T 23S rRNA gene mutation conferring linezolid resistance. Core genome phylogeny showed BX8117 was related to three other ST16 isolates from the UK, after removal of putative recombination blocks there were 76, 81, and 182 SNPs between these isolates suggesting they diverged from a common background but are not linked to recent transmission (Figure 2). ST16 has been associated with multidrug-resistant infections in humans and animals, highlighting the potential for the emergence of linezolid resistance in invasive enterococcal infections.^44^ The other five *optrA*-positive isolates have no close genetic links in this phylogeny (minimum pairwise SNPs 12,314 – 17,891). Our study is not designed to infer patterns across Scotland and the rest of the UK, but our findings suggest the *optrA*-positive isolates are generally distinct from those recently causing bloodstream infections in the UK.

**Figure 2.**
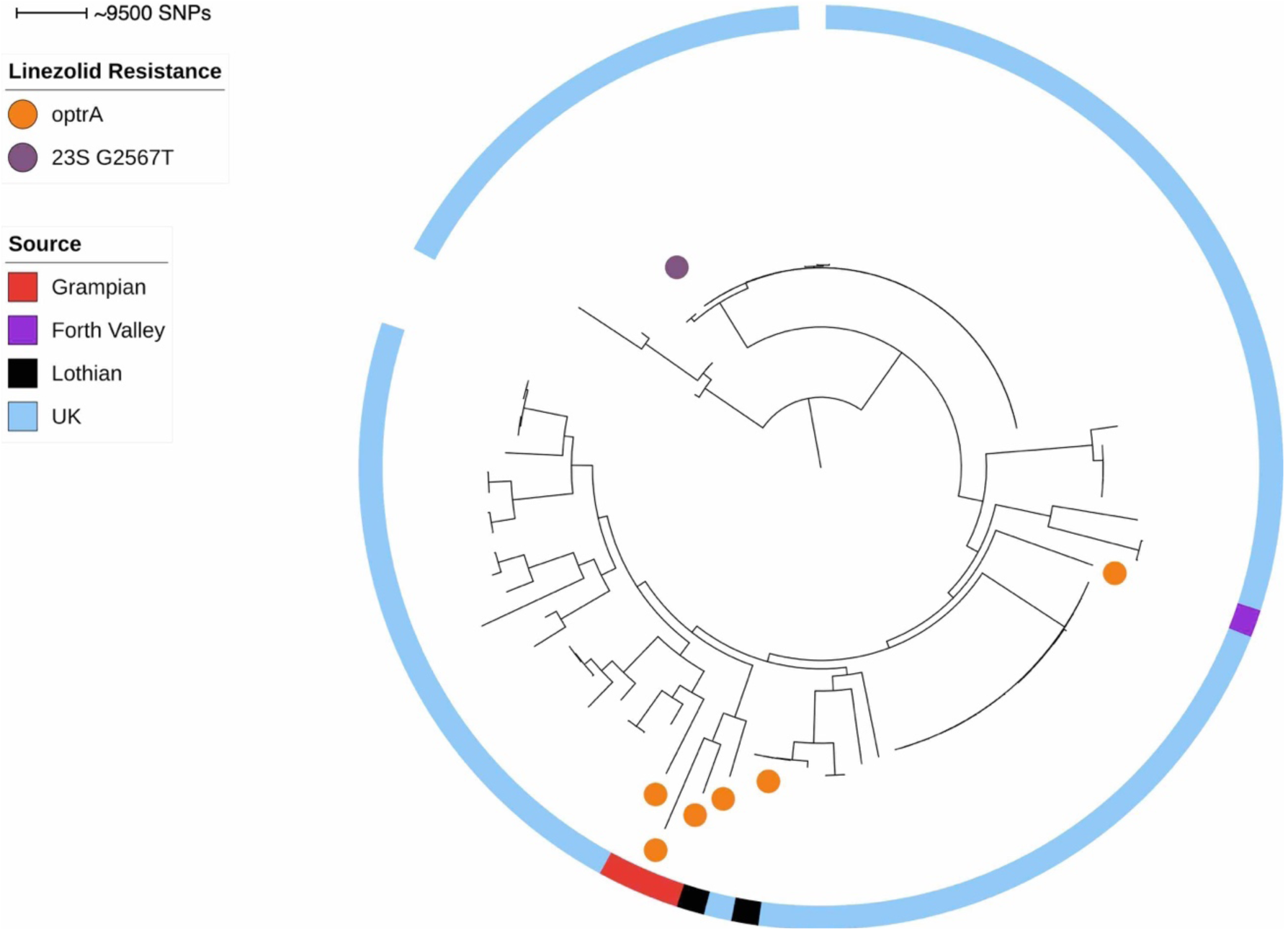
*optrA*-positive *E. faecalis* isolates in a national perspective. Phylogenetic analysis of the six *optrA*-positive isolates and 94 isolates from bloodstream infections in the UK shows the *optrA*-positive isolates are generally unrelated to others in the collection. Illumina reads were mapped to *E. faecalis* V583 reference genome, mobile genetic elements removed, and a maximum likelihood phylogeny performed on SNP alignment. Scale bar shows ∼9500 SNPs, linezolid-resistant isolates are indicated by circles, the outer ring indicates isolate source.

### *optrA*-positive *E. faecalis* harbour multiple resistance mechanisms

Looking at all assembled plasmids, the isolates carried genes conferring resistance to aminoglycosides (*ant*(6)-Ia, *aph*(3’)-IIIA, *aac*(6’)-Ie-*aph*(2’’)-Ia, and others listed in Table 2), chloramphenicol (*catA*8), bacitracin (*bcrA*), macrolides (*ermA*-like, *ermB*), tetracyclines (*tet*(L), *tet*(M)), trimethoprim (*dfrG*), and the heavy metals cadmium (*cadA*) and copper (*copZ*). However, the pattern of carried genes differed between isolates with only *optrA* and *fexA* found in all isolates (Table 2).

pBX8117-2 carried a gene with 100% nucleotide identity and coverage to *cfr*(D) from *Enterococcus faecium* isolated in France in 2015, and in Australia in 2019.^45,46^ In both isolates, *optrA* and *cfr*(D) genes were present on different contigs based on short-read sequencing. Our study is the first to detect *cfr*(D) in *E. faecalis* and using hybrid assembly we identified co-carriage of *optrA* and *cfr*(D) on the same plasmid. The French and Australian *cfr*(D)-positive isolates also carried *vanA*-type vancomycin resistance genes, although the Australian isolate was phenotypically vancomycin sensitive due to the loss of the regulatory genes *vanR* and *vanS*. At present, no *in vitro* work has described the impact of *cfr*(D) on antimicrobial resistance in enterococci so it is unclear whether or not *cfr*(D) confers the PhLOPS_A_ multiresistance phenotype originally described with Cfr.^47^

There is evidence of *optrA* being more common in particular *E. faecalis* lineages, with ST16, ST330, ST480, and ST585 in particular being described here and in other studies.^14,48–50^ These *optrA*-positive lineages are not specific to one host species and have been isolated from humans, animals, and the environment.^43,51^ Florfenicol use in food animals is associated with the presence of *optrA* in animal waste and the environment surrounding livestock farms.^52,53^ Additionally, *optrA*-positive enterococci and staphylococci have been isolated from raw food purchased from retail stores in China, Columbia, Denmark, and Tunisia.^51,54–56^ Wu *et al*. (2019) found evidence of transmission of *optrA*-positive *E. faecalis* from raw meat to a dog in China.^56^ However, the incidence of *optrA*-positive isolates in raw foods was low in the available studies, and there is currently no direct evidence to suggest *optrA*-positive strains are transmitted to humans via the food chain. ^57,58^ Increasing use of linezolid in human medicine may also select for *optrA-*positive strains, and once carried in the gut may be co-selected by other antimicrobials given the multidrug resistance phenotype of these isolates. The role of antimicrobial use, animal contact, food hygiene, and the environment in transmission of *optrA*-positive strains should be investigated further.

Our finding that *optrA* is present as different gene variants, carried on different mobile genetic elements, in unrelated strains of *E. faecalis* suggest a diverse *optrA* reservoir that is only partly investigated in this study. As well as *optrA*, the *cfr* and *poxtA* genes are emerging transferable linezolid resistance mechanisms. Further studies from a One Health perspective are warranted to understand the selection pressures driving transferable linezolid resistance, and the transmission dynamics of these strains to avoid further spread of linezolid resistance within *E. faecalis* and other Gram-positive bacteria.

## ACKNOWLEDGEMENTS

The authors would like to thank the Bioinformatics Unit at the University of St Andrews and Pathogen Informatics at the Wellcome Sanger Institute for access to high performance computing clusters.

## FUNDING

This work was supported by the Chief Scientist Office (Scotland) through the Scottish Healthcare Associated Infection Prevention Institute (Reference SIRN/10).

## TRANSPARENCY DECLARATION

The authors report no conflicts of interest related to this work.

---

**Figure S1.**
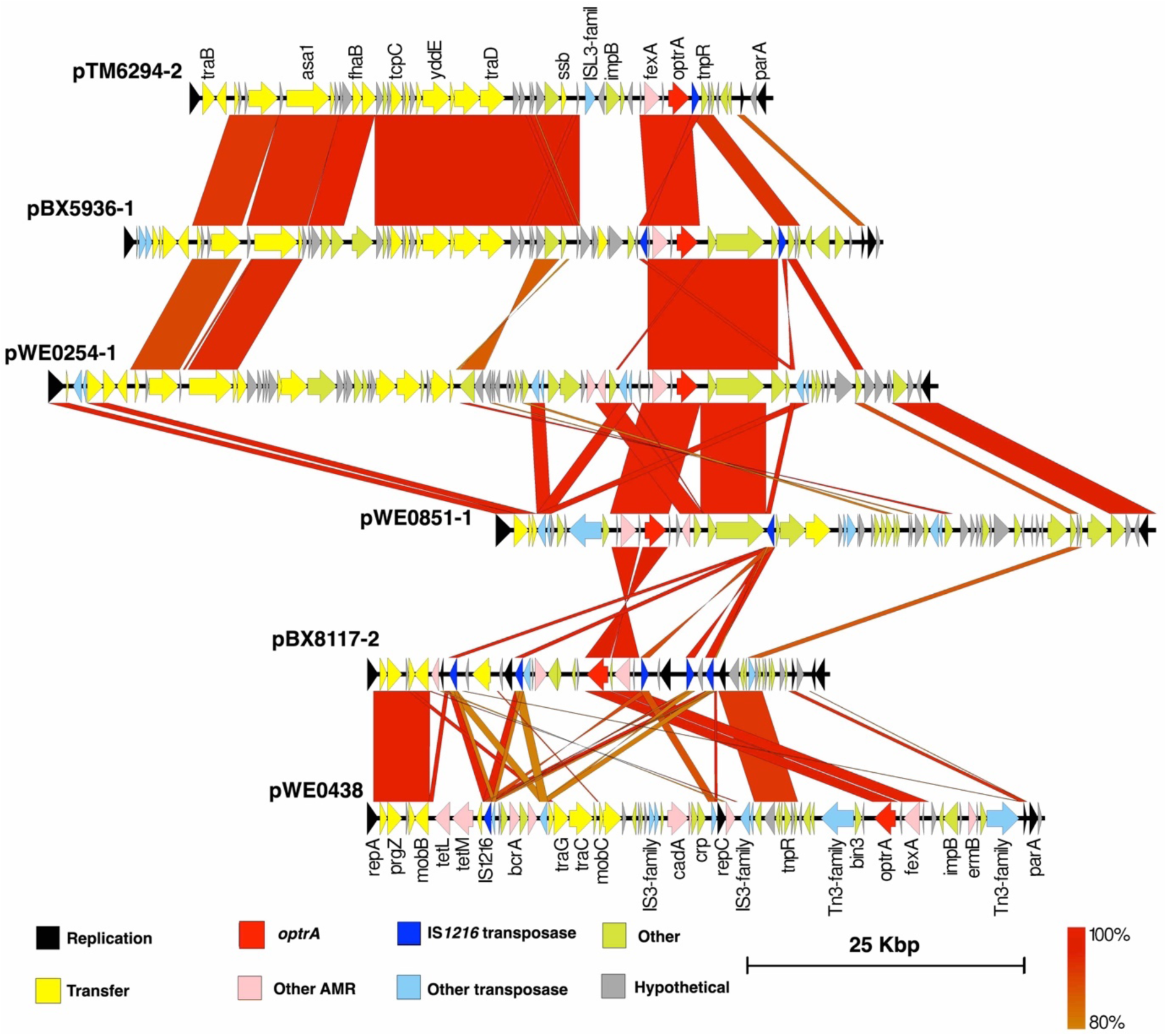
Alignment of full *optrA*-positive plasmid sequences. While some sequence similarity is seen between pTM6294-2 and pBX5936-1, in general identity is low between the *optrA*-positive plasmids, indicating *optrA* has mobilised to multiple plasmid backbones. Arrows indicate coding sequences, blocks between each sequence indicate regions with BLASTn sequence identity ≥80% and length >100bp.

**Figure S2.**
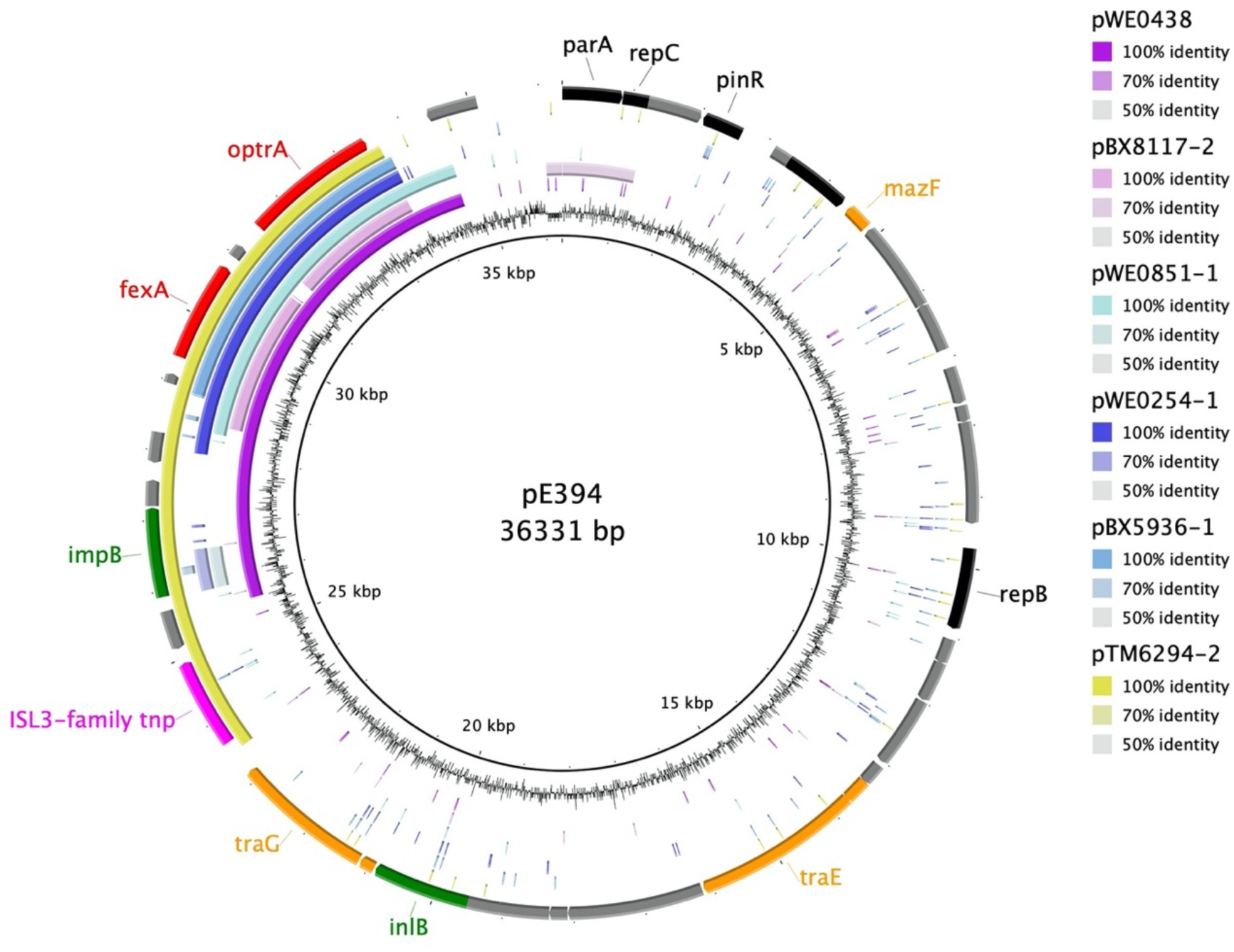
Alignment of full *optrA*-positive plasmid sequences against pE394. Sequence similarity confined to the *optrA*/*fexA* region. Inner ring indicates GC content of pE394, then alignment of pWE0438, pBX8117-2, pWE0851-1, pWE0254-1, pBX5936-1, pTM6294-2, and outer ring indicating coding sequences in pE394 (accession KP399637). Figure made with BRIG v0.95

